# Jasmonic acid and methyl jasmonate attenuate neuroinflammation via crosstalk with the prostaglandin E_2_/receptor EP2 signaling axis

**DOI:** 10.1101/2023.10.31.564983

**Authors:** Emily L Ward, Philip E Chen, Alaa Hussien-Ali

**Affiliations:** Royal Holloway, University of London, Department of Biomedical Sciences

**Keywords:** jasmonates, prostaglandin E_2_, neuroinflammation, EP2, neuroblastoma, neurodegeneration

## Abstract

The jasmonates are a class of oxylipin phytohormones known to exhibit anti-inflammatory, antioxidant, and anti-cancer effects in mammalian cells. We investigated the ability of three jasmonate compounds (jasmonic acid, methyl jasmonate, and 12-OPDA) and two structurally distinct jasmonate precursors (alpha-linolenic acid and palmitic acid) to attenuate inflammation in an *in vitro* model of neurodegenerative disease, for which the mechanisms of action have not been well identified. The study modeled chronic neuroinflammation in SH-SY5Y neuroblastoma cells using exogenous prostaglandin E_2_ (PGE_2_) treatment. Prostaglandin E_2_ caused concentration-dependent levels of inflammation and SH-SY5Y cell death, which were attenuated by the jasmonates and their precursors. To this end, structural similarities between the jasmonates and PGE_2_ were correlated with increased potency of their anti-inflammatory effects. Downstream biomarkers of signaling through the pro-inflammatory E prostanoid receptor subtype 2 (EP2) were then quantified using enzyme-linked immunosorbent assay methods. Of the compounds tested, only jasmonic acid and methyl jasmonate attenuated inflammation in the SH-SY5Y cells via crosstalk with the PGE_2_/EP2 signaling axis. Additionally, structural models and molecular binding simulations serve as evidence for our hypothesis that JA and MeJA achieve this crosstalk through competitive inhibition of the receptor EP2. This novel finding has implications in the study of neurodegenerative diseases for which the disease pathology is related to chronic neuroinflammation, including Alzheimer’s Disease (AD), Parkinson’s Disease (PD), amyotrophic lateral sclerosis (ALS) and multiple sclerosis (MS). In addition, these findings add to the understanding of the relationship between pro-inflammatory prostaglandin E_2_ signaling and disease severity.

## Introduction

The jasmonates (JAs) and prostaglandins (PGs) are classes of lipid-derived signaling molecules belonging to the oxylipin family, produced by the oxidation and reduction of polyunsaturated fatty acids from lipid membranes. Their ubiquitous presence and hormone-like signaling properties in plants and animals, respectively, suggest that the jasmonates and prostaglandins may be analogous between the plant and animal kingdoms. Across both kingdoms, these oxylipins respond to biotic and abiotic stressors by regulating various developmental, reproductive, and defensive processes on both the cellular and organismal scales (*Ahmad et al., 2016; Mueller, 1998; Wasternack, Song 2017*). These processes involve mechanisms that facilitate growth, fertility, and immune response in plants and animals. The ubiquitous synthesis and receptor-mediated, tissue-specific signaling of the oxylipins are collectively under the influence of developmental and environmental pressures, both endogenous and from outside injury or pathogens.

Plant jasmonates and mammalian prostaglandins are produced through biosynthesis pathways catalyzed by lipoxygenase (LOX) and cyclooxygenase (COX) enzymes, respectively. The LOX and COX enzymes are encoded by orthologous genes derived from an unknown ancestor common to plants and animals (*Kennedy 2014; Lee, Nioche, et al., 2008*). The oxidation and cyclization of membrane fatty acids through the LOX- and COX-associated pathways result in compounds that share a common partial cyclopentanone structure. Structural similarities between the jasmonates and prostaglandins are so significant that the jasmonates have proven useful skeletons for developing prostaglandin-like synthetic compounds. With the addition of enone functionality, jasmonate derivatives have prostaglandin-like anti-inflammatory properties and are more potent than naturally occurring prostaglandins themselves (*Dang et al., 2008)*.

In addition, naturally occurring plant jasmonates exhibit antioxidant, anti-inflammatory, and anti-cancer effects in both *in vitro* and *in vivo* models of human disease *(Alabi et al., 2019; Eduviere et al., 2015; Eduviere et al., 2016, Flescher, 2007; Goldin et al., 2008; McKenzie, Klegeris 2018; Taki-Nakano et al., 2016; Taki-Nakano et al., 2014).* However, the jasmonate mechanism of action in animals remains largely elusive. Our study aims to explore the mechanisms by which jasmonates might reduce neuroinflammation *(Taki-Nakano et al., 2016)* and lessen symptoms of neurodegenerative diseases in animals *(Eduviere et al., 2015)*, of which the pathology and disease progression are at least partially related to chronic neuroinflammation.

Extensive similarities in compound structure, function, and biosynthesis suggest that the cytoprotective activity of jasmonate signaling in mammalian cells might be attributed to crosstalk with that of endogenous mammalian prostaglandins.

Our study focuses on the use of 12-OPDA and two of its downstream derivatives, (+)-7-iso-jasmonic acid (jasmonic acid) and methyl-3 oxo-2-(2-pentenyl) cyclopentaneacetate (methyl jasmonate) (**Figure 1**). 12-OPDA, jasmonic acid (JA), and methyl jamsonate (MeJA) all share a partial cyclopentanone structure with prostaglandin E_2_. The production of trans jasmonic acid follows a reduction and 3 rounds of β-oxidation of cis 12-OPDA. The methylation of jasmonic acid then produces methyl jasmonate, altering its acetic acid functionality to an ester. JA and MeJA, but not 12-OPDA, thus share with PGE_2_ a trans configuration about the five-carbon ring. 12-OPDA and JA, but not MeJA, share the acetic acid moeity with PGE_2_.

**Figure 1.**
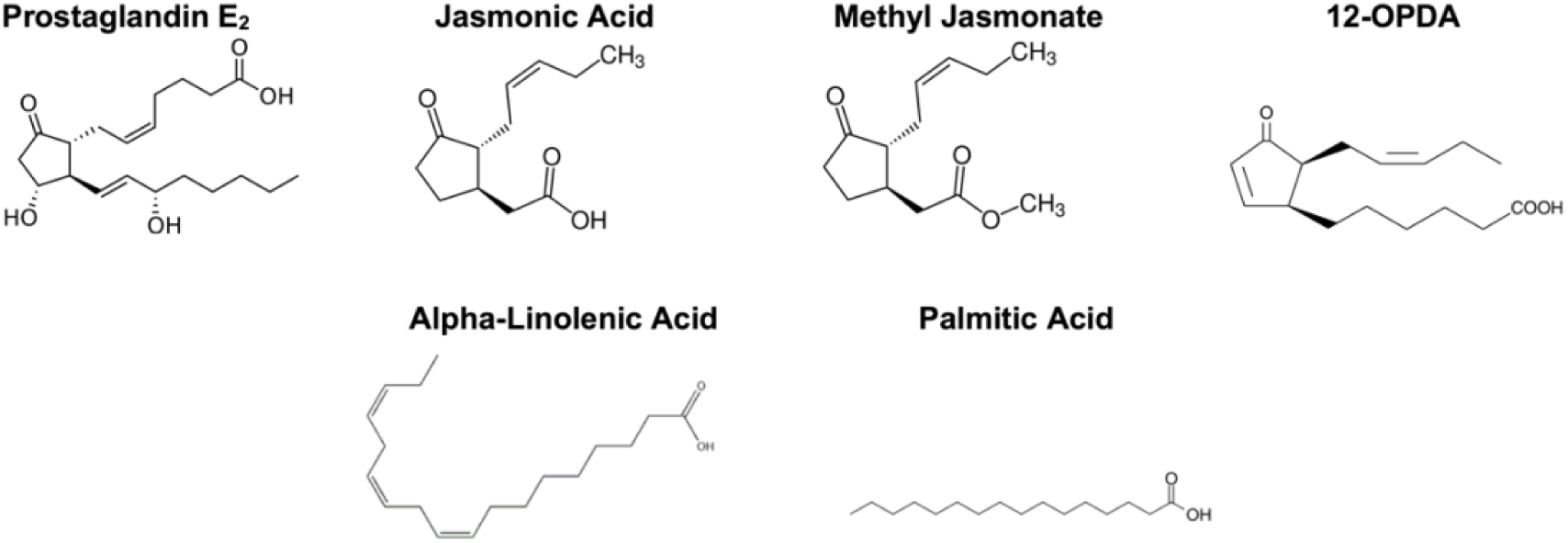
Structures of prostaglandin E_2_ and the jasmonates. Prostaglandin E_2_ shares a partial cyclopentanone structure with the jasmonates, including those used in the study (jasmonic acid, methyl jasmonate, and 12-OPDA). We used trans jasmonic acid and methyl jasmonate. 12-OPDA was in the cis configuration about the cyclopentanone ring. Methyl jasmonate and jasmonic acid share with PGE_2_ the acetic acid moiety, while 12-OPDA does not. The production of jasmonic acid follows a reduction and 3 rounds of β-oxidation of 12-OPDA. Methyl jasmonate is identical to jasmonic acid with the addition of a methyl group, altering its carboxylic acid functionality to an ester. Palmitic acid and alpha-linolenic acid are structurally distinct, bioactive precursors to the jasmonates.

**Figure 2.**
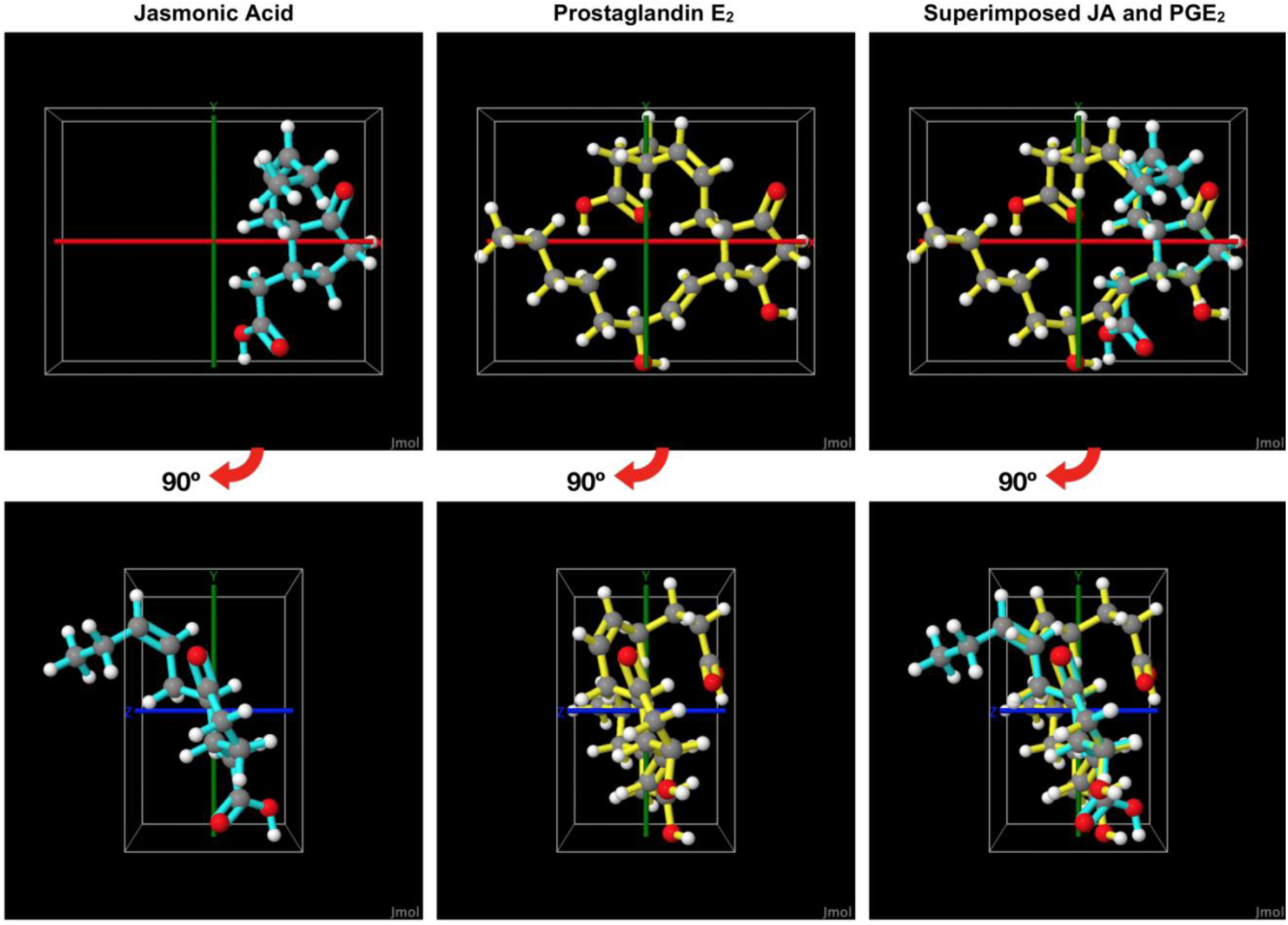
Jasmonic acid superimposed over prostaglandin E_2_. Of the three jasmonates, jasmonic acid most closely aligns with PGE_2_ when the two compounds are superimposed about their common cyclopentanone groups (RMSD= 1.68, 0.02 angstroms). JA and PGE_2_ share a partial cyclopentanone structure, the trans configuration about the ring, and the acetic acid moiety. For reference, the five-carbon ring and ketone functional groups can be seen along the positive X-axis of the top row. A 3-dimensional axis is included and has been set to a width of 0.1 angstroms for size reference. The bonds of the ball-and-stick model of jasmonic acid are cyan, and those of PGE_2_ are yellow. The below figures have been rotated 90 degrees from the top figures along the X-axis.

**Figure 3.**
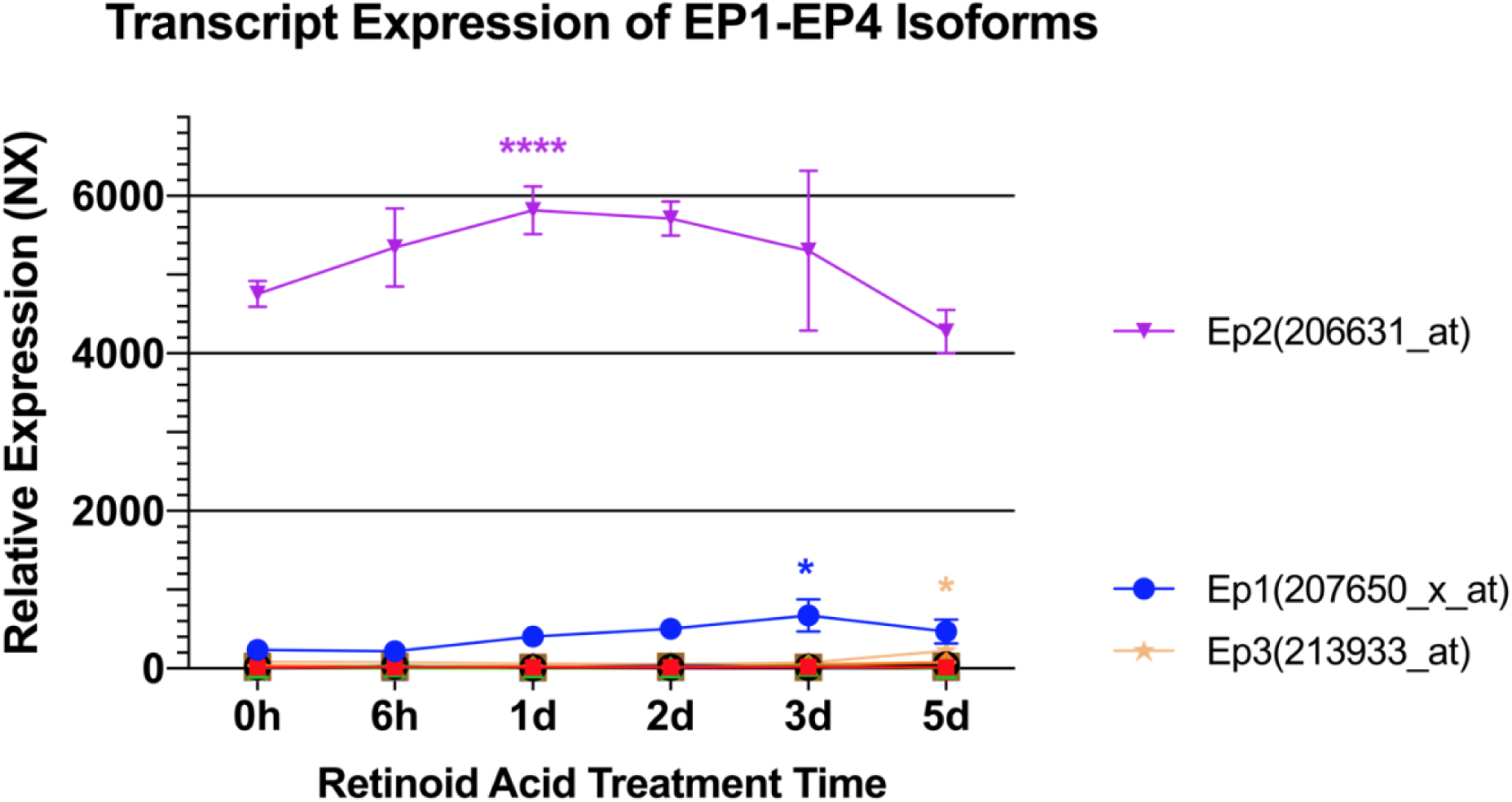
Using data from the paper *Nishida et al., 2008*, we analyzed the relative transcript expression of 16 isoforms of the E prostanoid receptor in SH-SY5Y cells treated with retinoic acid. Partial differentiation following 1 day of retinoic acid treatment increased EP2 (NCBI Protein accession number: 206631_at) transcript levels significantly by approximately 22%. Transcript levels of an EP1 isoform (207650_x_at) were significantly increased following 3 days of retinoic acid treatment, when transcript expression was increased by approximately 184% from day 0. The increases in transcript expression for both EP2 and EP1 were insignificant by day 5, when SH-SY5Y cells were considered terminally differentiated. One isoform of the receptor EP3 (213933_at) showed a significant rise of 165% in transcript expression following the full 5 days of retinoic acid treatment. These levels were still very low relative to the EP1 and EP2 receptor expression. All other EP isoforms lacked significant changes in transcript expression levels. A full list of the isoforms assayed for can be found in Supplemental Material.

We aim to assess whether jasmonate-modulated reductions in neuroinflammation proceed specifically via crosstalk with prostaglandin E_2_ (PGE_2_) signaling, a pivotal regulator of inflammatory network signaling in mammalian cells. In particular, we investigate activity through the prostaglandin receptor EP2 because of its role in modulating neuroinflammation and the innate immune response of the central nervous system *(Andreasson, 2010; Wang, 2019; Shi et al., 2010; Woodling, Wang, et al., 2014).* To this end, we measured inflammation in SH-SY5Y human neuroblastoma cells by monitoring PGE_2_-induced cell death. Novel treatment protocols were developed using this cell line and we observed jasmonate-attenuated inflammation indicated by increases in cell viability. 12-Oxo-phytodienoic acid (12-OPDA), jasmonic acid (JA), and methyl jasmonate (MeJA) reduced PGE_2_-induced inflammation and were associated with cytoprotective activity. Immunoassays for downstream biomarkers of receptor EP2 signaling indicated jasmonate-modulated reductions in inflammation were accompanied by decreased PGE_2_ signaling through the receptor EP2. We propose that reductions in PGE_2_/EP2 signaling associated with 12-OPDA and JA treatment might be attributed to direct competition for the EP2 receptor binding site, which could be the result of compound structure similarities. This theory was supported by the varying levels of response seen in the SH-SY5Y cells, which mirrored changes in the jasmonate treatment concentrations.

The results of this study add to the ongoing investigation of neuronal and microglial pathways associated with the cytoprotective effects of three of the most readily available and well-studied jasmonates: 12-oxo-phytodienoic acid, jasmonic acid and methyl jasmonate. Our findings also contribute to the current knowledge regarding the dual acting role of prostaglandin E_2_ in mediating inflammation through its various receptor subtypes. These findings have potential to assist in the discovery of biomarkers, drug targets, and therapeutic agents for the management of neuroinflammatory diseases.

## Materials and Methods

### Materials

Compounds used in this study are listed in Table 1. The jasmonates chosen are three of the most readily available and commonly studied. Two biosynthetic precursors to the jasmonates were also used (α-LA and PA) to assess the role of structural differences in the bioactivity of the jasmonates. Menadione was used as a control to induce measurable cell death. Alpha-tocopherol (Vitamin E), a known anti-inflammatory vitamin, was a control used to model recovery from PGE_2_-induced inflammation.

**Table 1.** Compounds used in this study. The compounds listed were used in either control experiments or as part of our primary investigation. Menadione was a control used to determine that we could reliably induce and measure concentration-dependent cell death in SH-SY5Y cells. A series of control pre-tests were then performed to ensure the compounds in question did not significantly alter untreated SH-SY5Y cell viability at the chosen concentrations. In our primary investigation of the jasmonates, alpha-tocopherol was a control compound that modeled recovery from PGE_2_-induced inflammation. Two jasmonate precursors (α-LA and PA) were used as structurally distinct controls to the jasmonates.

| Compound | Concentration | Primary Role |
| --- | --- | --- |
| Menadione | 5μM–50μM | (control) cytotoxicity |
| Prostaglandin E<sub>2</sub> (PGE<sub>2</sub>) | 30μM, 75μM | pro-inflammatory |
| α-Tocopherol (α-T) | 30μM | (control) anti-inflammatory |
| α-Linolenic Acid (ALA or α-LA) | 30μM | (control) jasmonate precursor |
| Palmitic acid (PA) | 30μM | (control) jasmonate precursor |
| Jasmonic acid (JA) | 30μM, 75μM | anti-inflammatory |
| Methyl jasmonate (MeJA) | 30μM | anti-inflammatory |
| 12-Oxo-phytodienoic acid (12-OPDA) | 30μM | anti-inflammatory |

Alpha-tocopherol, menadione, linolenic acid, palmitic acid, jasmonic acid, and methyl jasmonate were purchased from Sigma-Aldrich (St. Louis, MO, USA). Prostaglandin E_2_ was purchased from Tocris (Abingdon, UK). 12-OPDA was from Cayman Chemical (Ann Arbor, MI, USA). All compounds (10mM) were appropriately prepared as 10X working stocks using 100% ethanol as the solvent, with the exception of methyl jasmonate, for which double-distilled H_2_O was used.

Additionally, an enzyme-linked immunosorbent assay kit (UG_ADI-900-066) against intracellular cyclic-AMP was utilized from Enzo Life Sciences (St. Louis, MO, USA).

### Cell line culture

SH-SY5Y cells were obtained from our laboratory stock (originally purchased from the European Collection of Authenticated Cell Cultures) and were thawed down at the passage number of three. The SH-SY5Y cells were cultured with high glucose Dulbecco’s modified Eagle’s medium (DMEM) (Sigma-Aldrich) supplemented with 10% heat-inactivated fetal bovine serum (FBS) (Gibco/LifeTechnologies), 1% penicillin/streptomycin (10,000 U/ml, Gibco/LifeTechnologies), and 1% L-glutamine (200mM, Gibco/LifeTechnologies). We will refer to this media preparation as the 10% FBS DMEM media. Cell cultures were maintained in a 5.0% CO_2_ humidified atmosphere at 37 °C and split 1:5 at 80% confluency. To maintain homogenous and reproducible neuronal cultures, the epithelial-like cell phenotype of undifferentiated SH-SY5Y cells was maintained. This contrasts with the more expansive and branched neuron-like differentiated phenotype and is sufficient for our investigation of inflammatory signaling (*Shipley et al., 2016*). Undifferentiated SH-SY5Y cells may be kept for many weeks with regular passaging and cellular behavior, while partially differentiated and terminally differentiated SH-SY5Y cells display variable signaling effects and higher proportions of cell death when passaging (*Shipley et al., 2016)*.

To prevent differentiation, cells were cultured for ≤ 10 passages from when they were thawed (*Kawato et al, 2008; Shipley et al., 2016*). To this end, an advance in passage number was defined by the use of trypsin to split. We were careful also to observe the relative rate of cellular proliferation and cell morphology. That is, undifferentiated cells were expected to proliferate at a relatively rapid and constant rate, in contrast to the non-proliferating differentiated state. Cells were also expected to lack extensive outgrowth of neurites, or neuron-like extensions that seem to resemble dendrites and axons in the differentiated phenotype (*Kovalevich, Langford, 2016; Encinas et al, 2000*). This protocol offered a reliable and reproducible way to maintain the homogenous SH-SY5Y cell culture.

### Subculture of SH-SY5Y cells

A novel protocol was optimized for the treatment of the SH-SY5Y cells with prostaglandin E_2._ To this end, a different media preparation was required to elicit an inflammatory response in the cells. As per the advice of the authors of the paper *Miyagishi et al., 2013,* 1:1 DMEM/F-12 was completed with 0.5% FBS, 1% penicillin/streptomycin, and 1% L-glutamine. We will refer to this media preparation as the 0.5% FBS DMEM/F12 media. Cells were grown in a 5.0% CO_2_ humidified atmosphere at 37 °C and according to our original guidelines regarding passage, phenotype, and relative rate of proliferation. The 0.5% FBS DMEM/F12 media was used only in the plating and treatment of cells, but 10% FBS DMEM media was used for regular growth and passaging.

The 0.5% FBS DMEM/F12 media preparation was noticeably similar to the media used for the differentiation of SH-SY5Y cells, but it lacked the differentiating agent retinoic acid (*Shipley et al, 2016*). We inferred that SH-SY5Y cells required partial stress induced by serum starvation to respond to exogenous PGE_2_ with inflammatory signaling. This alternate cell culture requirement strengthens the notion that PGE_2_ signaling is subject to disease context.

### Cell viability assays

#### i. Dose-response tests

It was determined that we could reliably induce and measure concentration-dependent cell death amongst SH-SY5Y cells using a 48-hour incubation with 5-50µM menadione. Death was measured using trypan blue staining/visualization in combination with manual cell counting via haemocytometer. Data was displayed as a relative proportion of cell viability (%) in comparison to a vehicle control. The same protocol was used to determine whether treatment with the jasmonate compounds or their precursors significantly altered cell viability. The concentrations used were in the range of 15µM to 75µM. A target treatment concentration of 30µM was decided by consulting previous findings in the literature (*Taki-Nakano et al, 2014*; *Taki-Nakano et al, 2016*), and confirmed through these control pre-tests.

300µM working stocks of MeJA, JA, 12-OPDA, α-LA, PA and alpha-tocopherol were made using either ddH_2_O (methyl jasmonate only) or ethanol (all other compounds). 24 hours prior to treatment, SH-SY5Y cells were seeded in 6-well plates at a density of approximately 5.0 x 10^5^ cells/well. Following a 48-hour treatment incubation, a 100µL sample of cell suspension was taken from each well and stained using a 1:1 ratio with trypan blue. Each well was visualized and manually counted six times using haemocytometer slides loaded with 10µL of stained cell suspension. A singular count was represented as a proportion of cell viability, that is: *live cells / (stained cells + live cells*). Cell viability was assessed through a total of N=4 experiments per compound and the data from each experiment was normalized against the vehicle control before being displayed as relative proportions of viability.

#### ii. Prostaglandin E_2_-induced inflammation

SH-SY5Y cell death was induced with prostaglandin E_2_ according to a novel protocol that was optimized. 24 hours before treatment, cells were seeded at a density of approximately 2.5 x 10^5^ cells/well using 12-well plates. Cells were seeded using the alternate 0.5% FBS DMEM/F12 proliferation media, and then treated for 48 hours with 5µM-250µM PGE_2_ (see: ***Cell culture***). Cell viability was again assessed through trypan blue staining and manual haemocytometer counting (see: **Dose-response tests**). Because prostaglandin E_2_ increased the number of floating cells in each treatment well, both suspension and adherent cells were assessed for viability. This was performed for a total of N=4 experiments and data were analyzed and displayed identically to the control dose-response tests.

#### iii. Cotreatments with prostaglandin E_2_ and the jasmonates

Cells were again seeded in 12-well plates at a density of approximately 2.5 x 10^5^ cells/well using the 1:1 DMEM:F12 and 0.5% media (see: ***Cell culture***). After 24 hours, cells were treated using prostaglandin E_2_ alone or a combination of prostaglandin E_2_ and one of the jasmonates or precursors. The control was a vehicle control containing ethanol. We performed N=4 experiments for each prostaglandin E_2_ concentration: 30µM, 50µM, and 75µM. The jasmonate and precursor compounds were used at 30µM concentrations for each concentration of prostaglandin E_2_. We then examined a reverse response using 30µM prostaglandin E_2_ and increasing concentrations of jasmonic acid: 15µM, 30µM, and 75µM. After 48 hours, cell viability was again assessed using the same method previously described. This was performed for a total of N=4 experiments for each condition displayed in our results.

### PCR expression data analysis

Datasets from the Human Protein Atlas (HPA) (*Berglund et al., 2008*), NCBI Gene Expression Omnibus (GEO) (*Edgar et al., 2002*), and the primary study *Nishida et al., 2008* were cross-analyzed to examine the relationship between SH-SY5Y cell differentiation by retinoic acid and mRNA transcript level expression of genes encoding the prostaglandin E_2_ receptor subtypes EP1-EP4. Full access to raw data from the paper *Nishida et al., 2008* was found via the NCBI Gene Expression Omnibus (data accessible at NCBI GEO database (*Edgar et al., 2002*), accession *GSE9169*). From this we assessed the relative mRNA expression levels of 16 total isoforms of the E prostanoid receptors 1-4, and among two subtypes of SH-SY5Y cells (ECACC and ATCC). Included were three isoforms of *ptger1* transcripts, one of *ptger2*, nine of *ptger3*, and three of *ptger4.* A full list of the 16 isoforms included and their NCBI Protein accession numbers can be found in **Table A** of the Supplemental Material section.

Normalized expression of each mRNA isoform across treatment conditions was visualized using GEO online graphing. Comparisons between treatment conditions were visualized for all isoforms tangentially using GraphPad Prism for a total of N=3 datasets. A two-way ANOVA and Tukey’s multiple comparisons tests were performed to assess significance between the mean expression in the three experiments over time. Each treatment condition (6h, 1d, 2d, 3d, 5d) was analyzed for significance against the starting (0h) control. This was used to draw conclusions regarding prostanoid receptor expression amongst naïve, partially differentiated, and terminally differentiated SH-SY5Y cells. This information was used to make informed decisions regarding the optimization of our novel PGE_2_ treatment protocol.

### Enzyme-linked immunosorbent assays (ELISA)

SH-SY5Y cells were seeded in 12-well plates (2.5 x 10^5^ cells/ well) and treated for 48 hours with vehicle, PGE_2_ only (30µM or 75µM), or a cotreatments of PGE_2_ and one of the jasmonates (JA, MeJA, or 12-OPDA) or jasmonate precursors (PA or α-LA). Alpha-tocopherol was again used as a negative control against inflammation. These treatment conditions were identical to those used for the cell viability assays (see: ***Cotreatments with prostaglandin E_2_ and the jasmonates***). Suspension and adherent cells were lysed in 0.1M HCl for 10 minutes. The supernatants were assayed for intracellular cyclic-AMP using an enzyme-linked immunoassay kit (Enzo Life Sciences) according to the protocol of the manufacturer. A total of N=4 experiments were performed and strips of the 96-well microtiter plate (Enzo Life Sciences) were read using a microplate reader at 405nm (BioTek PowerWave 340).

Using 5 standards of various concentrations, as well as a blank, total activity (TA), NSB, and Bo condition in the microtiter plate, we created a 4PL standard curve in GraphPad Prism to interpolate mean cAMP concentrations (pmol/mL) from the optical densities (OD) recorded. A control test was performed to ensure that the assay detected rising cAMP levels using increasing concentrations of prostaglandin E_2_. The various cotreatment conditions were then assayed to examine the effect of jasmonate treatment on EP2 signaling, and cAMP levels were reported as pmol/mL.

### Compound structure analysis

Similarities in compound structure were assessed among prostaglandin E_2_ and each of the three jasmonates used. An image of the superimposition of prostaglandin E_2_ and jasmonic acid was created using Jmol: an open-source Java viewer for chemical structures in 3D (http://www.jmol.org/). A 360° rotation of this alignment was saved as a video using successive .jpeg images that differed 1° in viewing angle. Using the PubChem database (*Kim et al., 2019;*), prostaglandin E_2_ (compound ID number: 25280360) was loaded into Jmol and superimposed over jasmonic acid (CID: 7251180), methyl jasmonate (CID: 5281929), and 12-OPDA (CID: 5280411) individually. PGE_2_ and the jasmonates, which share a partial cyclopentanone structure, were aligned at the atoms of their five-carbon rings. The score of each alignment was used alongside further results of the study to make inferences about the importance of the shared partial structure in the efficacy of each jasmonate. The visualization aid was also used to assist in further molecular docking and the theoretical discussion of a proposed mechanism of jasmonate action.

### Molecular docking simulation

Jasmonic acid was docked into the E prostanoid receptor subtype 4 (EP4), the nearest structurally-related receptor to EP2 that exists as a file in the Protein Data Bank (*Breyer, 2001*). Jasmonic acid was chosen based on the results of the structural analysis performed (***see: compound structure analysis***), and the efficacy observed in our cotreatment tests for reduced inflammation (see: ***Results: Cotreatments with prostaglandin E_2_ and the jasmonates***). Using the jFATCAT-rigid algorithm from the Protein Data Bank, which reports domain-based structural alignments of proteins, we first confirmed significant structural similarities exist between EP2 and other relaxant cAMP-promoting receptors (DP, IP, and EP4) but not with constrictor receptors associated with decreased cAMP (EP1 and EP3). The results of this significance testing can be found in the Supplemental Material.

The Swissdock Molecular Docking program (*Grosdidier et al., 2011*) was used to upload a PDB file of EP4 and jasmonic acid as a ligand file from the ZINC compound database (*Irwin, Shoichet, 2006*). Files were opened and analyzed using UCSF Chimera software (*Petterson et al., 2004*), developed by the Resource for Biocomputing, Visualization, and Informatics at the University of California, San Francisco, with support from NIH P41-GM103311. The molecular docking was performed to strengthen the theoretical discussion of a jasmonate mechanism of action suggested by our findings.

### Statistical analysis

All preliminary data collected in our experiments was kept in Microsoft Excel. GraphPad Prism software (version 8.4.3, GraphPad Software Inc., La Jolla, CA, USA) was used to perform one-way ANOVA tests of significance, post-hoc multiple comparisons tests, and to create all graphs and figures. All primary cell viability data was normalized relative to the vehicle control (0µM) containing either ethanol or double distilled H_2_O, which was made equal to 100% viability. This included viability tests examining PGE_2_-induced inflammation. To assess the effects of cotreatment using prostaglandin E_2_ and the jasmonates, post-hoc comparisons were made between a PGE_2_-only treated control and each of the cotreatment conditions. Significance was established at P < 0.05. Cell viability data are represented as means ± standard error of the mean (SEM) unless otherwise stated.

The ELISA used to measure intracellular cAMP produced optical density data points for the conditions examined. These densities were transformed into interpolated concentrations of cAMP (pmol/mL) from a 4PL standard curve fit to the standard samples in the assay kit.

## Results

### Compound structure analysis

Jasmonic acid, methyl jasmonate, and 12-OPDA were superimposed over prostaglandin E_2_ using Jmol: an open-source Java viewer for chemical structures in 3D (http://www.jmol.org/). The jasmonates were aligned with PGE_2_ about their common partial structure, the cyclopentanone ring. Of the three jasmonates, jasmonic acid most closely aligned with PGE_2_ at the five-carbon ring. The root-mean-square-deviation of atomic positions (RMSD) was relatively low at 1.68, or 0.02 angstroms. Methyl jasmonate aligned similarly with a RMSD score of 1.85, also 0.02 angstroms. The loss of acetic acid functionality associated with the methylation of jasmonic acid (into methyl jasmonate) resulted in a slightly weakened alignment to PGE_2_.

12-OPDA did not superimpose well over PGE_2_ and the alignment resulted in a RMSD score of 2.03, or 1.03 angstroms. These results were to be expected as MeJA, JA, and PGE_2_ share a trans configuration about the cyclopentane ring, while 12-OPDA occurs in the cis configuration. The results suggest that both the trans configuration and acetic acid moiety are important factors to the structural alignment between jasmonates and PGE_2_, of which jasmonic acid has both.

The precursors alpha-linolenic acid and palmitic acid lack the same cyclopentanone ring and were not superimposed over PGE_2_. The jasmonate precursors were used as structurally unique controls that were likely to have similar cytoprotective effects (*Ali et al., 2019; Lee et al., 2018*). Distinctions in jasmonate and precursor activity were used to make inferences about the importance of the shared cyclopentanone ring to the jasmonate mechanism of action.

A 360° rotational video of the superimposition of jasmonic acid and prostaglandin E_2_ can be seen using the file listed with **Figure B** in the Supplemental Material.

### PCR expression data analysis

Using primary data reported in the paper *Nishida et al., 2008*, we analyzed the relative transcript expression of 16 isoforms of the E prostanoid receptor in SH-SY5Y cells treated with retinoic acid (**Figure 3**). The PCR expression data are plotted as means of N= 3 measurements with standard deviation (SD) included. In this instance, SD was used over SEM so error bars could be adequately visualized on the graph.

Of the sixteen isoforms of the E prostanoid receptor subtypes that were assayed, three isoforms displayed significant differences in transcript expression following some degree of retinoic acid treatment. Partial differentiation following 1 day of retinoic acid treatment increased EP2 (NCBI Protein accession number: 206631_at) transcript levels significantly (P < 0.0001), by approximately 22%. Transcript levels of an EP1 isoform (207650_x_at) were significantly increased following 3 days of retinoic acid treatment, when transcript expression was increased by approximately 184% from day 0 (P < 0.05). These increases in transcript expression for both EP2 and EP1 were attenuated and not significant by day 5, when SH-SY5Y cells were considered terminally differentiated.

One isoform of the receptor EP3 (213933_at) showed a significant rise (165%, P < 0.05) in relative transcript expression following the full 5 days of retinoic acid treatment. These levels were still very low relative to EP1 and EP2 receptor expression. Additionally, The Human Protein Atlas reports no normalized expression of the EP3 receptor (NX=0) in the SH-SY5Y cell line (*PTGER3: Cell Atlas (2015, March 12)*; *Berglund et al., 2008*). The disparity between our results and that of the HPA may be attributed to expression variances between other isoforms of EP3, of which there are eight (*Breyer et al., 2011)*, and the effects of normalizing the data. Still, in the partially differentiated state of SH-SY5Y cells that follows 1 day of retinoic acid treatment, the relative expression level of this EP3 isoform (NX=59.9) is nearly tenfold lower than that of the EP2 receptor (NX=5820.37). For this reason, we are not concerned that any expression of the anti-inflammatory receptor EP3 will significantly oppose the action of our target EP2 receptor. The other thirteen EP isoforms lacked significant changes in transcript expression levels. A full list of the isoforms assayed for can be found in **Table A** of the Supplemental Material section.

The results of our analysis suggest that terminal differentiation of the SH-SY5Y cells with retinoic acid would not significantly increase levels of either E prostanoid receptor expressed by SH-SY5Y cells, EP1 (NX= 4.1) or EP2 (NX= 10.2) (The Human Protein Atlas, *Berglund et al., 2008).* Rather, the data suggest a partially differentiated state might increase the expression of EP2 and promote an inflammatory response by SH-SY5Y cells to exogenous PGE_2_ treatment.

These results support the advice of the authors to the paper *Miyagishi et al., 2013* regarding the preparation of a media to be used when treating the cells with prostaglandin. As per the authors’ advice, it was decided that a differentiation-like (0.5% FBS) 1:1 DMEM:F12 media would be used. This media preparation was noticeably similar to the media used for SH-SY5Y cell differentiation, but the differentiating agent retinoic acid was not used.

We hypothesized that partial serum starvation caused by low levels of FBS might promote a partially differentiated cell phenotype that allows for prostaglandin E_2_ to significantly induced inflammation. This theory was supported by the literature, which confirms that stress signaling induced by gradual serum starvation is associated with progressive SH-SY5Y cell differentiation (*Shipley et al., 2016; Ricciotti, Fitzgerald, 2011*). In addition, further studies report that SH-SY5Y differentiation proceeds through cAMP-activated pathways (*Kume et al., 2008*), involving PKA and PI3K (*Sanchez et al., 2004*) and the ERK and p38 MAP Kinases (*Monaghan et al., 2008*). We therefore hypothesized that prostaglandin E_2_ might promote a positive feedback loop of stress signaling and antiproliferative pathways associated with chronic hyperinflammation.

In summary, the method of partial serum starvation was chosen in contrast to terminal differentiation with retinoic acid, which would take away from the maintenance of a homogenous cell line (see: ***Subculture of SH-SY5Y cells***) and which does not differ significantly in EP2 expression.

### Cell viability assays

Cell viability was examined as an indirect measure of cytotoxic inflammatory signaling in the SH-SY5Y cells (*Andreasson, 2010; Milatovic et al., 2011, Miyagishi et al., 2013*). That is, decreases in cell viability were indicative of increases in neurotoxic inflammation induced by PGE_2_. The anti-inflammatory vitamin alpha-tocopherol was used as a control to confirm that decreased inflammation was associated with improved cell viability.

#### i. Dose-response tests

We confirmed we could reliably induce and measure concentration-dependent cell death amongst SH-SY5Y cells using menadione. Following the same protocol, we monitored cell death in response to alpha-tocopherol, the jasmonates (jasmonic acid, methyl jasmonate, and 12-oxo phytodienoic acid), and two jasmonate precursors (alpha-linolenic acid, palmitic acid). The treatment concentration 30µM was taken from other primary studies using this cell line (*Taki-Nakano et al*, *2014*; *Taki-Nakano et al, 2016*), and has no significant effect on SH-SY5Y cell viability for any of the compounds after 48 hours of treatment (see: **Supplemental Material** for figures).

#### ii. Prostaglandin E_2_–induced inflammation

A novel protocol was created to induce inflammation in SH-SY5Y cells using prostaglandin E_2_. The protocol used the 0.5% FBS DMEM/F12 media preparation and treatment with 5µM-250µM PGE_2_ for 48 hours. Inflammation was indicated by reduced cell viability and recorded as a percentage of cell survival, relative to the vehicle control. At concentrations of 20µM and higher, PGE_2_ treatment was associated with significantly decreased cell viability. PGE_2_ induced cell death in a concentration-dependent manner in the supraphysiological treatment range of 20µM to 250µM. Our results agreed with those using similar cell lines (*Miyagishi et al., 2013*).

Cells supplemented with the 10% FBS DMEM proliferation media, and thus void of serum starvation, did not respond to exogenous PGE_2_ treatment. PGE_2_ treatment using 10% FBS DMEM did not significantly influence cell viability in comparison to the vehicle control, indicating that the cells were void of either inflammatory or anti-inflammatory signaling induced by exogenous PGE_2_. These results suggest that SH-SY5Y cells must be under partial stress to respond to PGE_2,_ as is supported by the results of the section ***PCR expression data analysis*** and an analysis of the relevant literature (*Kume et al., 2008*; *Miyagishi et al., 2013*; *Monaghan et al., 2008; Shipley et al., 2016; Sanchez et al., 2004).* Partial serum starvation accomplished using the 0.5% FBS DMEM/F12 media allowed for an inflammatory response by PGE_2_. These results support the notion that PGE_2_/EP2 signaling responds to disease context and contributes to neuronal death after inflammation has progressed from acute to chronic (*Andreasson, 2010; Lima et al., 2012; Milatovic et al., 2011; Wang et al., 2019*).

#### iii. Cotreatments with prostaglandin E_2_ and the jasmonates

To investigate whether the jasmonates attenuate PGE_2_-induced inflammation in the SH-SY5Y cells, the next round of experiments used cotreatments of prostaglandin and each of the jasmonates. The precursors α-LA and PA were also included, bioactive jasmonate precursors that lack the same partial cyclopentanone structure to PGE_2_, JA, MeJA, and 12-OPDA. Alpha-tocopherol was used as a control known to reduce inflammation and promote cell viability. Cell viability relative to a vehicle control was again used as an indirect measure of cytotoxic inflammation induced by prostaglandin E_2_ treatment.

As expected, the next round of experiments displayed significant cell death induced by 30µM PGE_2_ (**Figure 5. A**). This was indicated by a strong level of significance (P < 0.0001) in a post-hoc comparison of the prostaglandin-only treated control (PGE_2_) and the vehicle control (C). The relative percentage of cell survival at 30µM PGE_2_ (34.20%, **Figure 5. A**) was notably lower than in our previous investigation of prostaglandin–induced inflammation alone (47.15%, **Figure 4**). Still, we consider the 30µM treatment condition a successful control as it fell within the expected range indicated by our first round of experiments, i.e. between 27.19% survival (50µM PGE_2_, **Figure 4**) and 65.80% survival (20µM PGE_2_, **Figure 4**). This discrepancy in survival rates at 30µM PGE_2_ might have been reduced by an increased number of tests in either set of experiments.

**Figure 4.**
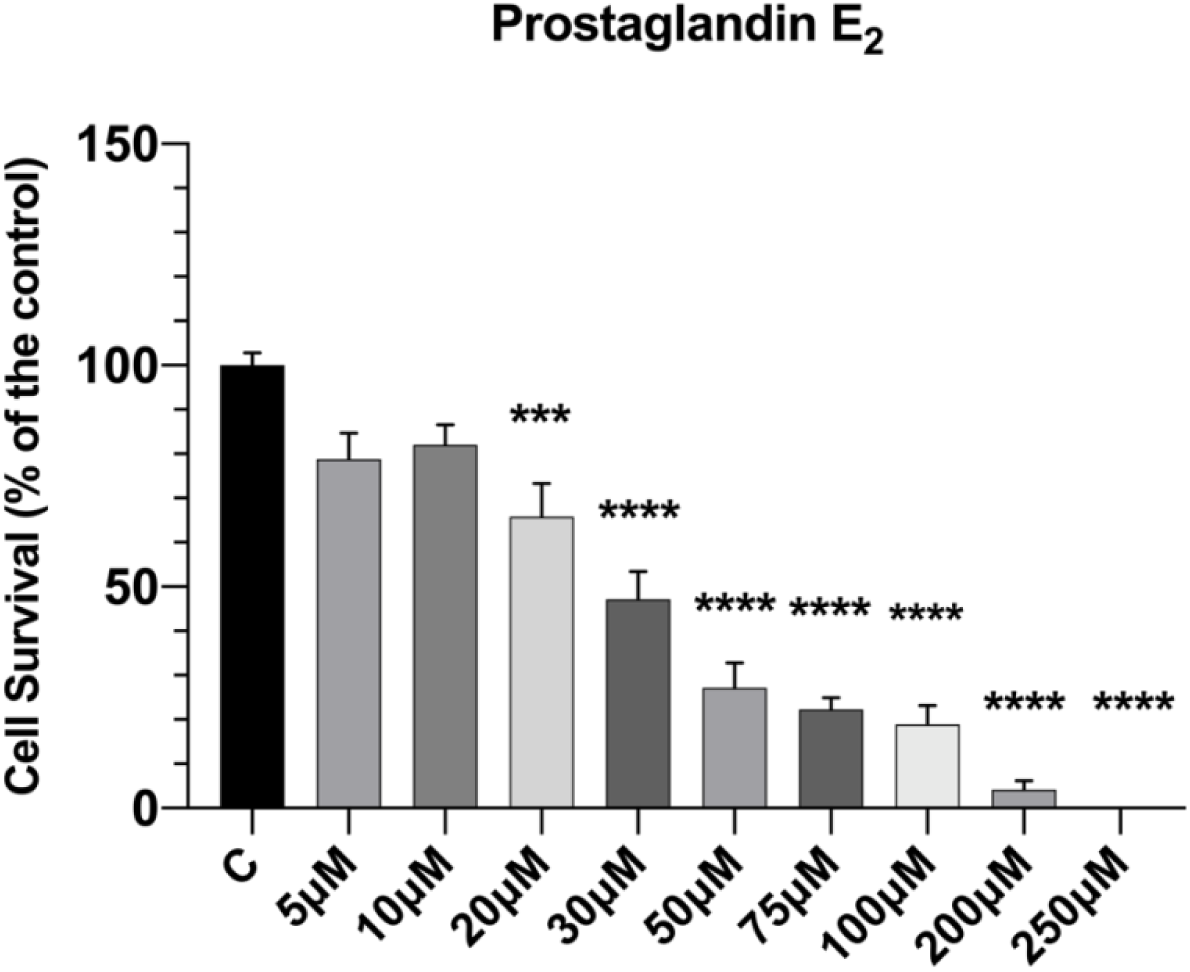
Treatment with exogenous prostaglandin E_2_ causes significant inflammation associated with SH-SY5Y cell death in the range of 20µM to 250µM. Cell death occurs in a concentration-dependent manner following 48-hours of treatment.

**Figure 5.**
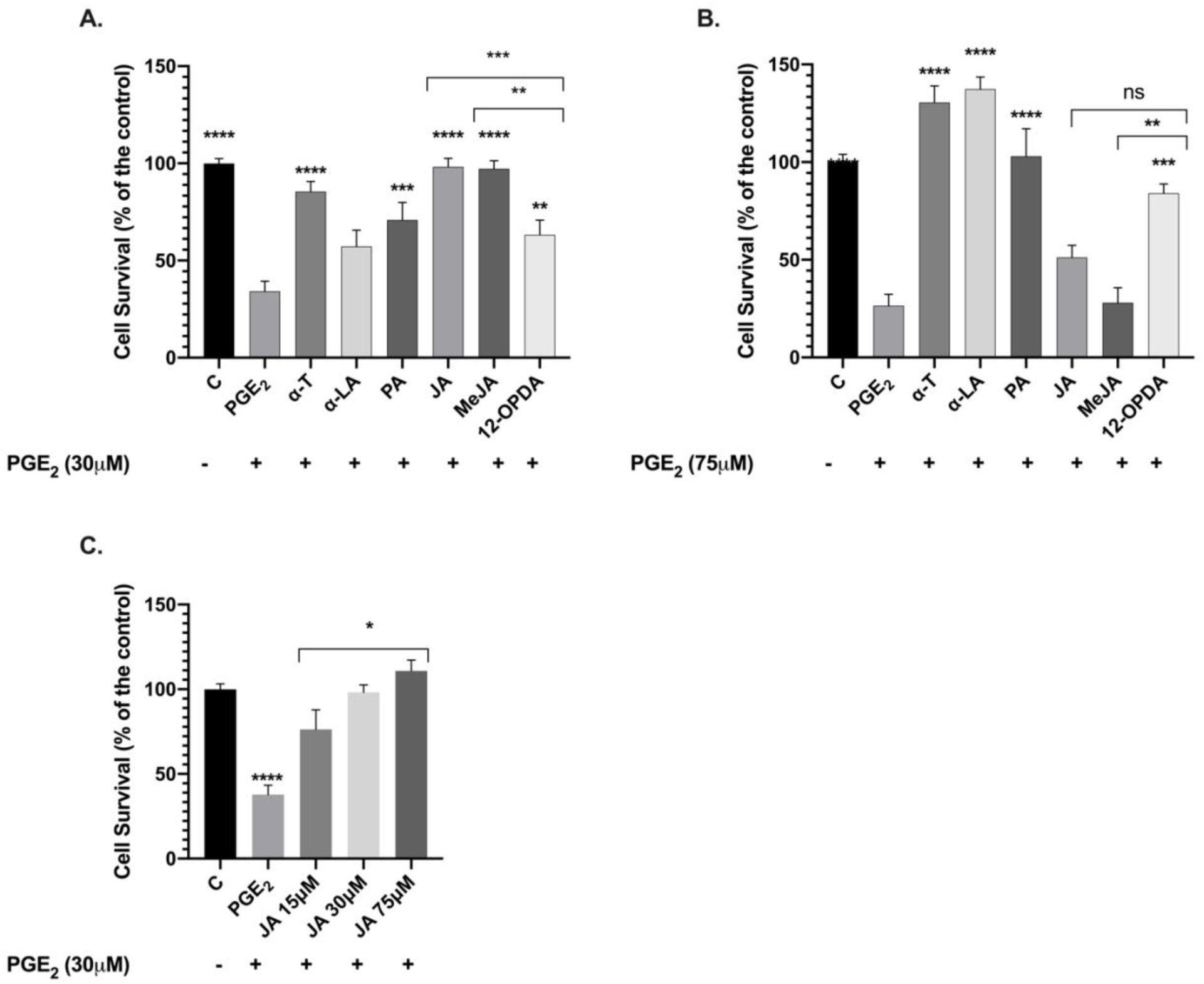
(**A**) JA, MeJA (P < 0.0001),12-OPDA (P < 0.01), and PA (P < 0.001) significantly attenuate SH-SY5Y cell death by opposing inflammation induced with 30µM PGE_2_. The jasmonates most structurally similar to PGE_2_ (i.e. JA and MeJA) were most potent in their anti-inflammatory effects. The effects of JA and MeJA did not significantly differ. The extent to which JA and MeJA promoted cell viability exceeded the potent anti-inflammatory control alpha-tocopherol (α-T). (**B**) 12-OPDA (P < 0.001), α-LA, and PA (P < 0.0001) significantly attenuate inflammation and cell death induced by 70µM PGE_2_. Alpha-tocopherol and α-LA appear to have elevated cell viability above the vehicle control, but this observation was not statistically significant. (**C**) Jasmonic acid ameliorates 30µM PGE_2_-induced inflammation and cell death in dose-dependent manner. The anti-inflammatory effects of 75µM jasmonic acid significantly exceeded those of 15µM jasmonic acid.

SH-SY5Y cell death induced with 30µM prostaglandin E_2_ was significantly attenuated by 30µM alpha-tocopherol (P < 0.0001), the anti-inflammatory control compound (**Figure 5. A**). 30µM jasmonic acid (JA), methyl jasmonate (MeJA), 12-OPDA, and palmitic acid (PA) also significantly attenuated the cell death associated with 30µM PGE_2_ treatment. Alpha-linolenic acid (α-LA) was associated with increased cell viability, but this did not differ significantly from the control treated with PGE_2_ only. Jasmonic acid and methyl jasmonate potently promoted cell viability above even the anti-inflammatory control alpha-tocopherol (α-T). JA and MeJA also increased viability to nearly 100% of the vehicle control, representing near-complete recovery from 30µM PGE_2_-induced inflammation. The anti-inflammatory effects of 12-OPDA were significantly less potent than those of JA and MeJA, though greater than either of the precursor compounds.

Our findings indicate that structural differences amongst the jasmonates and their precursors may be related to the potency of their anti-inflammatory effects. In the presence of neuroinflammation induced by 30µM PGE_2_ treatment, jasmonates containing a cyclopentanone ring, and the trans configuration about the ring, displayed a greater ability to attenuate inflammation than those that are in the cis configuration or lack the partial cyclopentanone structure. The acetic acid moiety did not appear to be related to the efficacy of the jasmonates, as jasmonic acid and methyl jasmonate did not differ significantly in the extent to which they attenuated cell death.

Following the experiments involving 30µM jasmonate treatments and 30µM PGE_2_, we investigated the anti-inflammatory capabilities of 30µM jasmonate treatments in the presence of 75µM PGE_2_ (**Figure 5. B**). We altered this ratio between the jasmonate and prostaglandin treatment concentrations in order to assess our hypothesis that the jasmonates may directly interfere with the binding of prostaglandin to the receptor EP2. We predicted that the jasmonates would be less effective at reducing inflammation, i.e. at promoting cell viability, when the concentration of PGE_2_ was increased.

As expected, relative cell survival was lower after treatment with 75µM prostaglandin E_2_ (26.49%, **Figure 5. B**) than it was after using 30µM prostaglandin E_2_ in the prior experiments (34.20%, **Figure 5. A**). The proportion of cell survival closely mirrored that reported in our initial experiments using 75µM prostaglandin E_2_ to display prostaglandin–induced inflammation (22.32%, **Figure 4**). As predicted by our hypothesis, 30µM treatments of JA and MeJA no longer significantly attenuated PGE_2_-induced inflammation (**Figure 5. B**). Interestingly, alpha-linolenic acid, palmitic acid, and 12-OPDA did significantly promote cell viability despite a greater concentration of prostaglandin used. 12-ODA, α-LA, and PA all displayed increased efficacy as the PGE_2_ concentration was increased. Further, cell viability associated with α-LA treatment did not differ significantly from that of alpha-tocopherol (the anti-inflammatory control) or palmitic acid. Under treatment with 75µM prostaglandin E_2_, alpha-tocopherol and α-LA appear to have elevated cell viability above the vehicle control, but this observation was not statistically significant.

Taken together with the previous findings, these results suggest that the anti-inflammatory compounds tested (α-LA, PA, JA, MeJA, and 12-OPDA) are all capable of attenuating neuroinflammation induced by prostaglandin E_2_ in SH-SY5Y cells. The potency at which the compounds reduce neuroinflammation appears to be related to compound structure and the relative level of inflammation present, or the concentration of prostaglandin E_2_ used. Those compounds most similar in structure to PGE_2_ (i.e. JA and MeJA) seem to act through a different mechanism than those that are structurally distinct (12-OPDA, α-LA, and PA). At lower concentrations of PGE_2_, the mechanism of action orchestrated by JA and MeJA is more effective at reducing inflammation. At higher PGE_2_ concentrations, there may be different anti-inflammatory mechanisms allowing for greater efficacy of 12-OPDA, α-LA, and PA. In part, these results might be explained by the known ability of PGE_2_ to respond to disease severity by activating different intracellular signaling pathways (*Andreasson, 2010; Lima et al., 2012; Milatovic et al., 2011; Wang et al., 2019*). It follows that activity through the PGE_2_/EP2 signaling axis differs across varying levels of neuroinflammation (*Wang et al., 2019*). If the jasmonates attenuate inflammation through crosstalk with the PGE_2_/EP2 signaling axis, then perhaps this happens at different levels in the signaling pathway depending on the compound structure.

We then aimed to strengthen this argument by again using cell viability as a measure of 30µM PGE_2_-induced inflammation. Cotreatments using prostaglandin and varying concentrations of jasmonic acid were chosen in search of a reverse effect to that already seen, which might further suggest competition for the receptor EP2. Jasmonic acid was chosen due to its structural similarities to PGE_2_ (see: ***Compound structure analysis***) and the potency of its anti-inflammatory effects at 30µM PGE_2_ (**Figure 4)**. Additionally, among the control pretests performed (see results in **Supplemental Material, Figure A**), only JA lacked significant changes in cell viability when used alone at a treatment concentration of 75µM.

Jasmonic acid attenuated the inflammation induced by 30µM prostaglandin E_2_ in a concentration-dependent manner, indicated by increases in relative cell viability (**Figure 5. C**). In post-hoc comparisons to the vehicle control, none of the jasmonic acid treatment conditions (15µM, 30µM, or 75µM) varied significantly in cell viability. These results indicated that, at concentrations of 15µM and higher, jasmonic acid allowed for complete recovery of 30µM PGE_2_–induced inflammation in the SH-SY5Y cells. 75µM jasmonic acid was significantly more potent in its ability to promote cell viability at 75µM than at 15µM (P < 0.05). These results were to be expected if jasmonic acid and prostaglandin E_2_ were competing for binding of the pro-inflammatory receptor EP2. To assess this further, we performed enzyme immunoassays and measured activity through the EP2 receptor in each of these conditions.

### Enzyme-linked immunosorbent assays (ELISA)

Enzyme-linked immunosorbent assays were used to measure intracellular levels of cyclic-AMP, a pro-inflammatory second messenger associated with the PGE_2_/EP2 signaling axis. After prostaglandin E_2_ binds and activates the GPCR receptor EP2, the G-protein-mediated activation of adenylyl cyclase leads to the formation of cAMP. cAMP-activated pathways promote inflammation, and thus, cAMP is often used as a measure of inflammatory signaling in SH-SY5Y cells (*Kang et al., 2017*).

Assays were carried out to assess whether the anti-inflammatory effects of the jasmonates were associated with reduced prostaglandin E_2_ signaling through the receptor EP2. To this end, we hypothesized that JA and MeJA might compete with prostaglandin E_2_ binding to EP2 and result in decreased levels of cAMP in the presence of 30µM PGE_2_. Recorded cAMP concentrations were normalized versus the prostaglandin-only treated control in each experiment and post-hoc comparisons were performed using these normalized values (see: **Supplemental Material**, **Figure C**). Each calculated significance level was then transferred to the graphs containing discrete concentrations of cAMP that were averages of N=4 experiments. The anti-inflammatory control alpha-tocopherol was not included in this set of experiments, as it reduces inflammation through a mechanism that does not include interaction with the PGE_2_/EP2 signaling axis (*Devaraj et al., 1996; Lee et al., 2006*).

Control tests confirmed that levels of cAMP detected in SH-SY5Y cells increased significantly with rising PGE_2_ treatment concentrations (**Figure 6. A**). Cells treated with a vehicle control measured a mean baseline cAMP concentration of 0.193 pmol/mL (SEM ± 0.025) and that number nearly doubled to 0.385 pmol/mL (SEM ± 0.043) following 48 hours of treatment with 200µM PGE_2_. These results indicated that the previous inflammatory PGE_2_ signaling we observed proceeded through the receptor EP2. We confirmed that prostaglandin treatment is associated with inflammation mediated by EP2 signaling, which results in rising cAMP and eventual cell death through the activation of cAMP-related pathways. Increased cAMP levels were associated with cytotoxicity, and improved cell viability was expected to be correlated with decreased cAMP.

**Figure 6.**
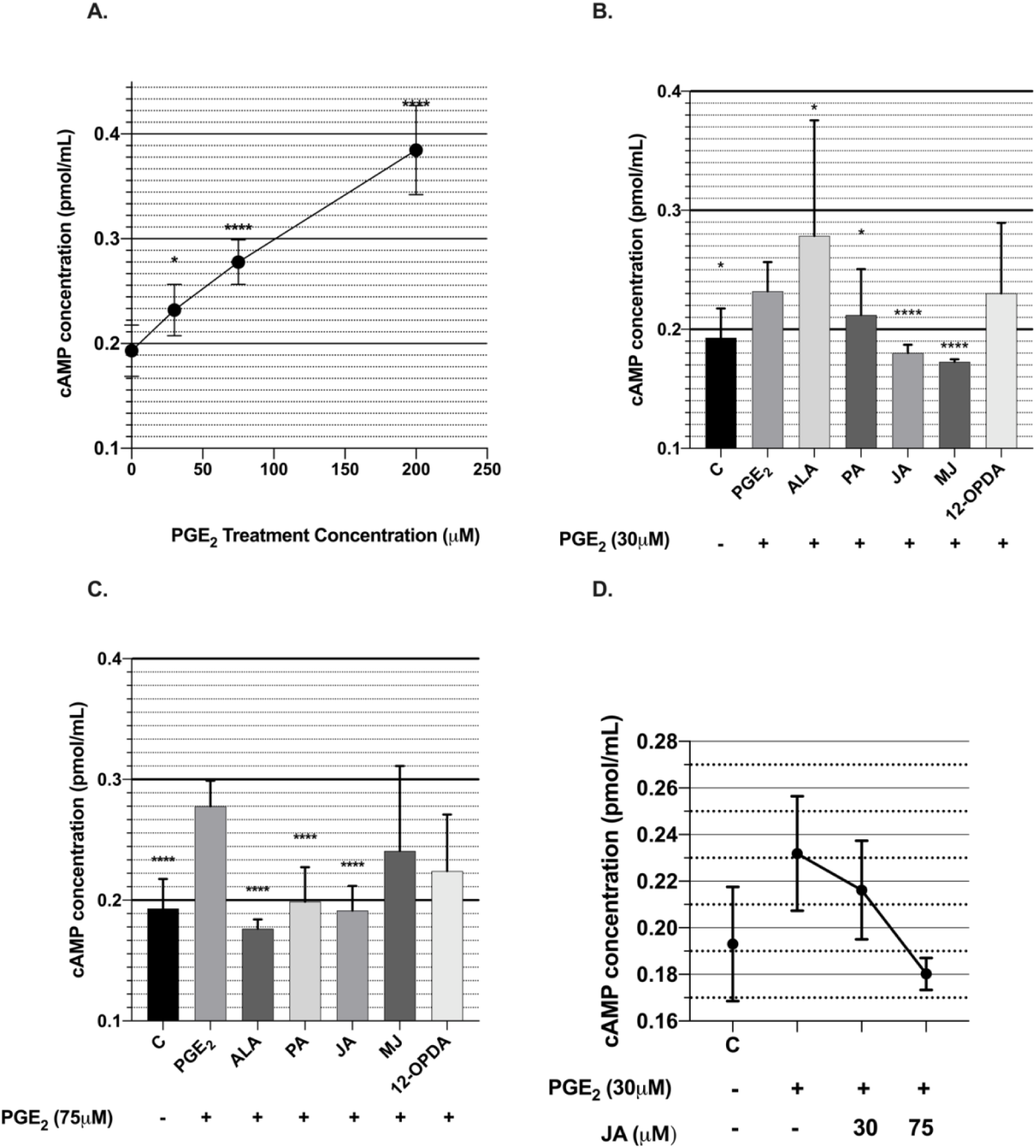
(**A**) Increasing treatment concentrations of PGE_2_ were positively correlated with intracellular cAMP levels in SH-SY5Y cells (measured in pmol/mL). Cells treated with a vehicle control measured a mean baseline cAMP concentration of 0.193 pmol/mL (SEM ± 0.025), which nearly doubled to 0.385 pmol/mL (SEM ± 0.043) following 48 hours of treatment with 200µM PGE_2_. (**B**) Jasmonic acid (JA) and methyl jasmonate (MeJA) most significantly attenuated increased cAMP levels associated with 30µM PGE_2_ treatment. cAMP levels associated with 30µM α-LA were significantly higher than in cells treated with PGE_2_ alone. Post-hoc comparisons were made relative to the PGE_2_-only treated control. (**C**) Jasmonic acid (JA), alpha linolenic acid (α-LA), and palmitic acid (PA) significantly attenuated increased cAMP levels associated with 75µM PGE_2_ treatment (P < 0.0001). Methyl jasmonate and 12-OPDA did not significantly lower cAMP levels. The standard error associated with MeJA (SEM ± 0.07036) and 12-OPDA (SEM ± 0.04688) was relatively high. (**D**) The mean intracellular concentration of cAMP associated with 30µM PGE_2_ treatment was significantly reduced in the presence of jasmonic acid. This reduction in cAMP responds to increasing concentrations of JA.

Next, cells cotreated with 30µM PGE_2_ and the jasmonates were assayed for intracellular cAMP. The results of the immunoassay were assessed through post-hoc comparisons made relative to the PGE_2_-only treated control. The results mirrored those seen in our viability assays for relative inflammation **(Figure 5. A**), in which methyl jasmonate and jasmonic acid most significantly opposed the actions of 30µM PGE_2._ Enzyme immunoassays reported that jasmonic acid (JA) and methyl jasmonate (MeJA) most significantly attenuated the increased cAMP levels associated with 30µM PGE_2_ treatment (**Figure 6. B**). These findings indicate that MeJA and JA are associated with decreased PGE_2_ signaling through the EP2 receptor. In contrast, alpha-linolenic acid, palmitic acid, and 12-OPDA all did this to a lesser extent or not significantly at all. These results were also consistent with those seen in the previous set of viability experiments. Interestingly, cAMP levels associated with 30µM α-LA were significantly higher than in cells treated with PGE_2_ alone. Therefore, these results suggest that the jasmonic acid and methyl jasmonate mechanisms of action interfere with the PGE_2_/EP2 signaling axis in the presence of 30µM PGE_2_. However, the results do not strongly support a mechanism of action orchestrated by α-LA, PA, or 12-OPDA that relies on crosstalk with the PGE_2_/EP2 axis. Rather, our results suggest together that the structurally distinct compounds to PGE_2_ (α-LA, PA, and 12-OPDA) are associated with different signaling mechanisms than MeJa and JA, and that these mechanisms seem variable across inflammatory conditions (i.e. PGE_2_ concentration).

In the next set of immunoassays, performed using 75µM PGE_2_, cAMP levels were significantly elevated above a vehicle control as expected **(Figure 6. C**). Intracellular cAMP was also higher at 75µM PGE_2_ than in the previous experiments using 30µM PGE_2_. In line with the results of our viability assays (**Figure 5. B**), palmitic acid and alpha-linolenic acid potently reduced the inflammatory effects of 75µM prostaglandin E_2_. However, the effects of the jasmonates at this PGE_2_ concentration did somewhat differ from those seen in the viability assays. That is, 12-OPDA did not significantly attenuate the increase in cAMP caused by PGE_2_ treatment, though it significantly promoted cell viability. Additionally, jasmonic acid significantly reduced cAMP while it did not significantly improve cell viability. These results affirm that structural differences amongst the jasmonates contribute to alternate mechanisms of signaling responsible for their array of cytoprotective effects. Specifically, this set of immunoassays provides evidence that jasmonic acid most potently interacts with the PGE_2_/EP2 signaling axis at both concentrations of PGE_2_, and that this crosstalk is enough to promote SH-SY5Y viability in the presence of 30µM PGE_2_ but not 75µM PGE_2_. Methyl jasmonate appears less potent in its ability to interrupt EP2 signaling or to promote cell viability. These results are therefore first evidence that the potencies at which MeJA and JA interact with the PGE_2_/EP2 signaling axis may differ. Because jasmonic acid can be more closely aligned to PGE_2_ than MeJA, we believe this could be evidence that crosstalk occurs in the form of competitive inhibition for the EP2 receptor. Additionally, 12-OPDA appears to reduce inflammation through a separate mechanism that does not reduce activity through the EP2 receptor. The remaining compounds appear also to act through distinct mechanisms that vary across intracellular context and result in realtively large error between experiments.

To further assess the ability of jasmonic acid to interrupt the PGE_2_/EP2 signaling pathway, we again tested the effects of increasing jasmonic acid concentrations in the presence of 30µM PGE_2_ (**Figure 6. D**). Under increasing concentrations of jasmonic acid, cAMP levels in PGE_2_-treated SH-SY5Y cells decreased significantly in a concentration-dependent manner. These results suggest that the level of crosstalk between jasmonate and prostaglandin signaling, and thus the level of jasmonate bioactivity, is a function of their concentration in SH-SY5Y cells. These results support a theory of direct competition for binding of the EP2 receptor. Intertest variability in cAMP concentration was high, but data normalized against the control for each experiment was used to test significance. Intertest variability resulted in high ranges of error, but the effects seen in the comparison of group means were mirrored within each test. The high rate of intertest variability could have been due to disparities in cell density, though we put forward our best effort to keep the cell culture homogenous and consistent. In this case of error, greater cell density would heavily influence cAMP concentrations upwards.

### Molecular docking simulation

We simulated the molecular docking of jasmonic acid into the prostaglandin E_2_ receptor EP4 to assess the potential for JA to compete with PGE_2_ for binding with the prostaglandin receptor. The simulation produced a best-fit model in which the functional groups of jasmonic acid were positioned closely to the amino acid residues indicated in the binding of prostaglandin E_2_ to the receptor (**Figure 7. A, B**). The study *Toyoda et al., 2019* reported that the carboxylic acid group of prostaglandin E_2_ interacts with the guanidinium group of Arginine 316 and the hydroxyl group of Tyrosine 80 to form hydrogen bonds with the EP4 receptor. The carboxylic acid group of PGE_2_ also forms hydrogen bonds with Threonine 168. These residues can be seen positioned closely to the carboxylic acid group of jasmonic acid in the best-fit model produced by SwissDock.

**Figure 7.**
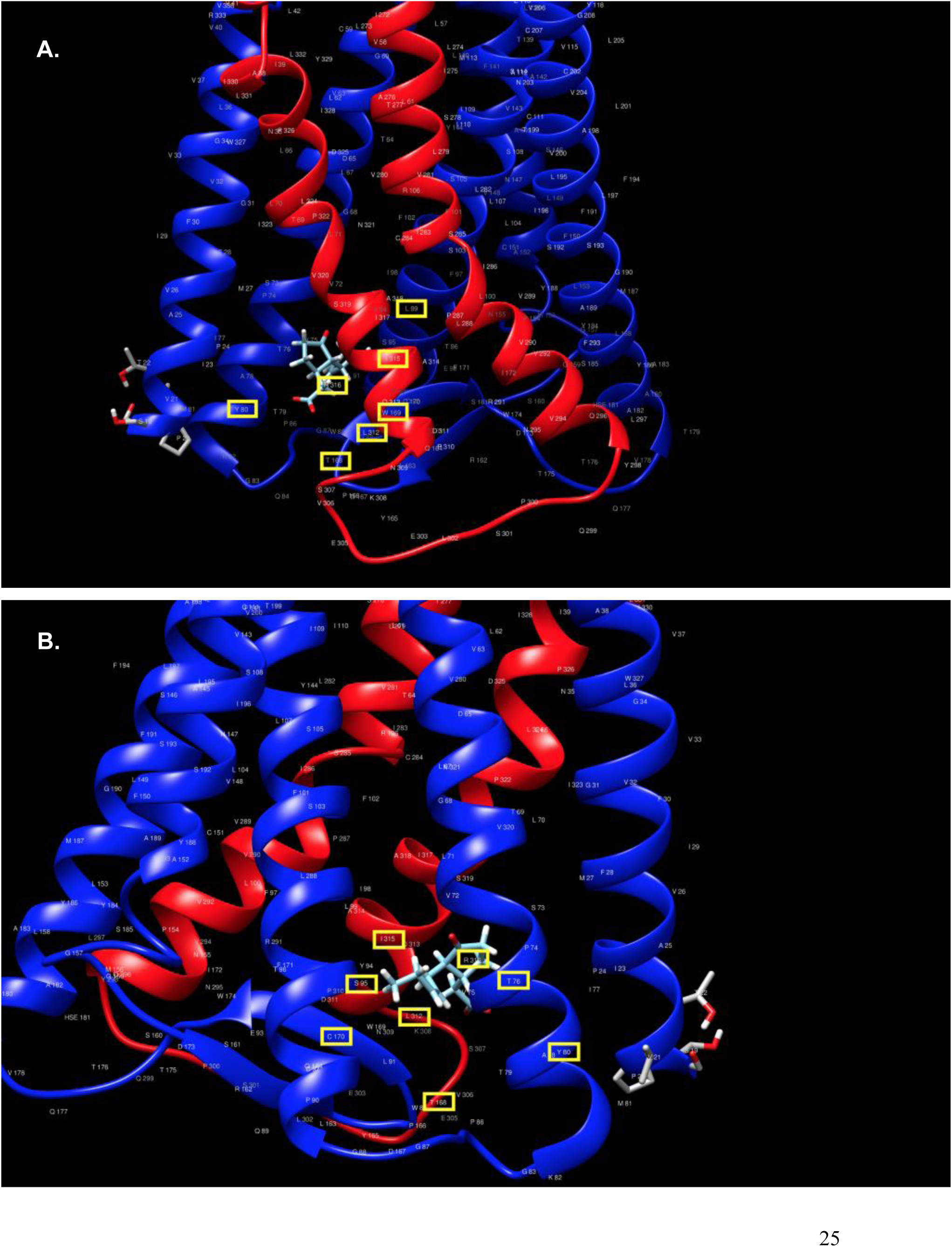
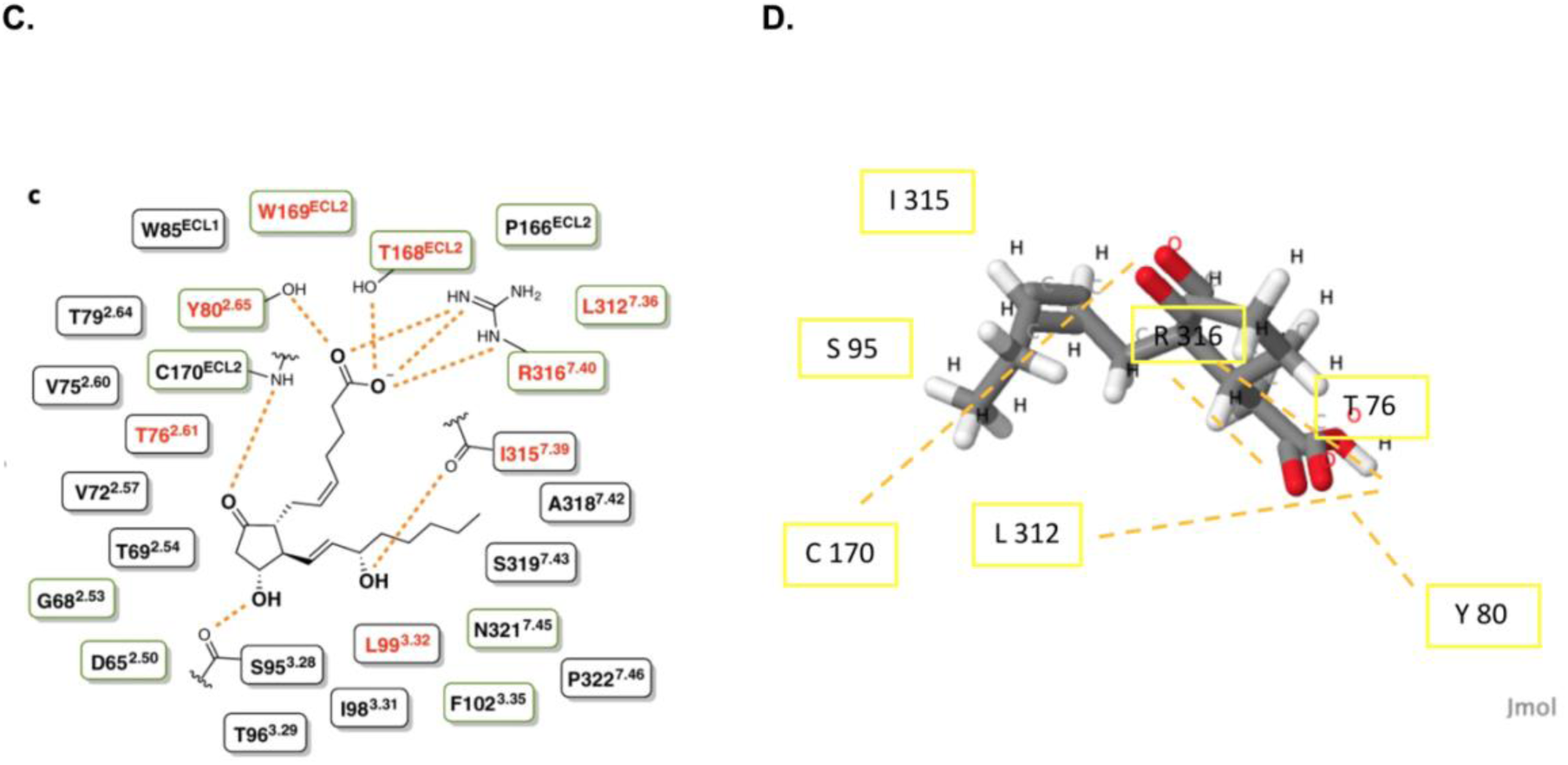
(**A**) The molecular docking of jasmonic acid into the receptor EP4. The residues essential to the binding of prostaglandin E_2_ into the receptor have been indicated (*Toyoda et al., 2019*). (**B**) A 90-degree rotation from Figure A. Residues essential to PGE_2_ binding have again been indicated. (**C**) Taken from the paper *Toyoda et al., 2019,* indicating the hydrogen bonds formed between PGE_2_ and the essential residues of EP4. These residues have been indicated in Figures A and B. (**D**) Indicating the potential hydrogen bonds formed between functional groups of jasmonic acid and amino acid residues within the binding pocket of EP4. The results were made using structural similarities between JA and PGE_2_, an analysis of the hydrogen bond donors and acceptors in the binding of PGE_2_.

The backbone amide of Cysteine 170, the carbonyl group of Serine 95, and carbonyl group of Isoleucine 315 form an additional three hydrogen bonds to the 9-oxo,11-hydroxyl, and 15-hydroxyl groups of PGE_2_, respectively. The study also reports that Threonine 76, Leucine 99, Tryptophan 169, and Leucine 312 are important to the binding of PGE_2_ to the EP4 receptor (**Figure 7. C**). These residues in the prostaglandin binding pocket are also indicated in the images of the molecular docking of jasmonic acid into the receptor. Shared functional groups between jasmonic acid and PGE2 seem to position themselves near the same essential amino acid residues, suggesting potential for molecular binding by hydrogen bonds.

An analysis of structural similarities between jasmonic acid and prostaglandin E_2_, in addition to the observed binding orientation and proximity score (Δ= −8.65 kcal/mol with a FullFitness of - 1048.73) reported by Swissdock, supports the hypothesis that jasmonic acid may be capable of binding to the similar receptor EP2. Pictured is the top-scoring docking pose of jasmonic acid in EP4. This hypothesis supports the theory that the jasmonate mechanism of action is, at least in part, related to direct competition for the prostaglandin receptor EP2.

In addition, we include a side-by-side representation of the binding interactions *possible* in the docking of each ligand (it is important to note that jasmonic acid may not bind to the receptor, in which case these interactions are chemically plausible but may not occur) (**Figure 7. D**). These results indicate that JA may be similar enough in structure to prostaglandin E_2_ to cause competitive inhibition of the pro-inflammatory EP2 receptor. However, further biochemical studies are needed to confirm these potential binding interactions. At current, a Protein Data Bank file does not exist for the receptor EP2. Future studies may focus on this receptor that appears to be strongly indicated in chronic inflammation and neurodegenerative disease.

## Discussion

This *in vitro* study established a novel protocol for inducing inflammation and cell death in SH-SY5Y neuroblastoma cells using treatment with supraphysiological levels of prostaglandin E_2_. Prostaglandin E_2_ (20µM to 250µM) caused significant cell death associated with hyperinflammation in a concentration-dependent manner. The cells, which express only the EP1 and EP2 receptor subtypes, were assayed for cAMP as a downstream indicator of PGE_2_/EP2 axis signaling. An enzyme-linked immunosorbent assay against intracellular cyclic-AMP confirmed that the inflammatory signaling induced by prostaglandin E_2_ in the cells was associated with signaling through the receptor subtype EP2. That is, increased treatment concentrations of PGE_2_ were associated with increased levels of intracellular cAMP in the SH-SY5Y cells. It was presumed that subsequent cell death was caused by cAMP-activated pathways (*Takadera et al., 2004*) and the cumulative cytotoxicity of pro-inflammatory species signaling. The study modeled chronic inflammation and neurodegenerative disease, during which excess inflammatory response signaling orchestrated by prostaglandin E_2_ is responsible for the production of cytotoxic inflammatory cytokines, ROS, and RNS that contribute to neuronal cell death (*Andreasson, 2010; Wang et al., 2019*).

The standard *in vivo* concentration of PGE_2_ in neurons is not known, but spinal cord levels of pro-inflammatory PGE_2_ in models of neurodegenerative disease have been reported around 150 pg/mg tissue (0.42µM) (*Klivenyi et al., 2004*). Other studies report that PGE_2_ signaling through the receptor EP2 is anti-inflammatory at concentrations less than 10nM (*Chen et al., 2018*). Our results add evidence to the theory that the effects of PGE_2_/EP2 axis signaling change in response to pathophysiological context *in vivo*, i.e. disease severity, and with PGE_2_ concentration *in vitro* (A*ndreasson, 2010; Wang, 2019; Shi et al., 2010; Woodling, Wang, et al., 2014*). As such, we add to the ongoing investigation of the mechanisms by which prostaglandin E_2_ signaling can be pro-inflammatory, anti-inflammatory, cytotoxic, and cytoprotective, etc., all within a given process.

Our study modeled changes in pathophysiological context using two preparations of proliferation media for our SH-SY5Y cell culture. In the presence of normal 10% FBS DMEM media, SH-SY5Y cell viability is unaffected by supraphysiological levels of PGE_2_ treatment. In contrast, SH-SY5Y cells experiencing partial serum starvation due to the use of 0.5% FBS DMEM/F12 media display significant hyperinflammation and cell death in response to PGE_2_. These observations mirrored those in the literature regarding the progressive degeneration of neurons across advancing stages of chronic neuroinflammation and neurodegenerative disease (*Andreasson et al., 2010; Innamorato et al., 2008*). The results support the theory that PGE_2_ signaling drives chronic inflammation, contributing to cytotoxicity and neuronal death, when the immune response is chronically activated (*Milatovic et al., 2011*).

Due to the role of chronic inflammation in the progression of neurodegenerative disease, we sought to examine the therapeutic potential of a class of anti-inflammatory phytohormones, the jasmonates. The naturally occurring plant jasmonates exhibit antioxidant, anti-inflammatory, and anti-cancer effects in both *in vitro* and *in vivo* models of human disease, and are remarkably similar in structure and function to mammalian prostaglandins *(Alabi et al., 2019; Eduviere et al., 2015; Eduviere et al., 2016, Flescher, 2007; Goldin et al., 2008; McKenzie, Klegeris 2018; Taki-Nakano et al., 2016; Taki-Nakano et al., 2014).* We hypothesized that the jasmonates may reduce inflammation via crosstalk with the PGE_2_/EP2 signaling axis, particularly in the form of competitive inhibition for the EP2 receptor. We theorized that competition for EP2 binding may exist if structurally similar compounds to PGE_2_ ameliorate inflammation more potently than those with greater structural differences.

Three jasmonate compounds (jasmonic acid, methyl jasmonate, and 12-OPDA) and two structurally distinct jasmonate precursors (alpha-linolenic acid and palmitic acid) were therefore assessed for their potential to modulate PGE_2_-induced inflammation in the neuroblastoma cells. An assessment of compound structure similarities and molecular docking simulations elucidated the potential for JA and MeJA to compete for the EP2 receptor, while 12-OPDA, α-LA, and PA appeared significantly distinct in structure to PGE_2_. The anti-inflammatory potential of each compound was assessed through cotreatments with PGE_2_ and cell viability assays, in which cell death was an indirect measure of cytotoxic inflammation.

Cell death induced by prostaglandin E_2_ was attenuated by the jasmonates and their precursors, but the potency at which each compound reduced inflammation was related to its structural similarities with PGE_2_. To this end, differences in compound structure seemed to indicate different cytoprotective mechanisms that varied in effectiveness across PGE_2_ concentration. Jasmonic acid and methyl jasmonate, which share a partial cyclopentanone structure and trans configuration with PGE_2_, were not distinct in their cytoprotective effects but were significantly different in potency to the effects of the cis compound 12-OPDA and the structurally distinct precursors α-LA and PA. That is, at 30µM PGE_2_ treatment, JA and MeJA were most effective at promoting cell viability. In the presence of 75µM PGE_2_, the three structurally distinct compounds (12-OPDA, α-LA, and PA) were more effective at reducing cell death.

Further enzyme-linked immunosorbent assays for intracellular cyclic-AMP suggested that JA and MeJA act through the same anti-inflammatory mechanism: crosstalk that results in decreased signaling of the PGE_2_/EP2 signaling axis. The acetic acid moiety shared by jasmonic acid and PGE_2_ also seemed to increase the potency at which jasmonic acid reduced EP2 signaling in comparison to MeJA. Further, increasing concentrations of jasmonic acid ameliorated PGE_2_-induced cell death and reduced levels of cAMP in a concentration-specific manner, suggesting competition for the receptor EP2.

In comparison, when cotreatments of PGE_2_ with 12-OPDA, palmitic acid, and α-LA were assayed for intracellular cAMP, the variable results produced high amounts of error and showed little consistency to the results of the previous viability assays. These discrepancies suggest the mechanism of action orchestrated by these structurally unique compounds is likely different to that of JA and MeJA and doesn’t rely on crosstalk with the PGE_2_/EP2 signaling axis. As the jasmonates are a family of phytohormones with an array of cytoprotective properties, we propose that this array of actions may be possible due to small structural changes that strongly influence compound signaling and downstream effects. This theory is in agreement with observations of the mammalian prostaglandins, which achieve an array of biological activities through small structural differences.

The results of this study add to the incomplete knowledge surrounding the jasmonates and their mechanisms of action in mammalian cells. Additionally, we provide results and protocols to be used to examine the dual role of prostaglandin E_2_ signaling in the context of neurodegenerative disease. Our experiments reliably and measurably reduced neuroinflammation in SH-SY5Y neuroblastoma cells, representing promise in the study of an important aspect of neurodegenerative disease pathologies that include Alzheimer’s Disease, Parkinson’s Disease, amyotrophic lateral sclerosis and multiple sclerosis. Ultimately, the jasmonates pose therapeutic potential in the management of inflammatory diseases and the promotion of cell viability both *in vitro* and *in vivo*. Future studies might further examine the potential for JA or MeJA to interact with the mammalian receptor EP2 through biochemical methods. Additionally, we suggest future projects use neuronal/glial cocultures to explore potential of the microglial receptor EP4 due to the ability of microglia to alter SH-SY5Y neuroinflammation (*Pandur et al., 2018*).

## Acknowledgements/ Conflicts of Interest

The authors declare no conflicts of interest.

