## Supplemental Figures for "Jasmonic acid and methyl jasmonate attenuate neuroinflammation via crosstalk with the prostaglandin E_2_/receptor EP2 signaling axis"

**Figure A. Dose-response tests performed as controls** Treatment with menadione induces cell death in concentration-dependent manner (**A**). It is confirmed that we can detect and measure concentration-dependent cell death in SH-SY5Y cells using this protocol and staining with trypan blue. Neither the compounds used as control measures (**B,C,G**) nor the jasmonate compounds in question (**D-F**) significantly alter SH-SY5Y cell viability at treatment concentrations 30μM. The data compiles n=4 experiments using each compound ang is displayed as means ± SEM.

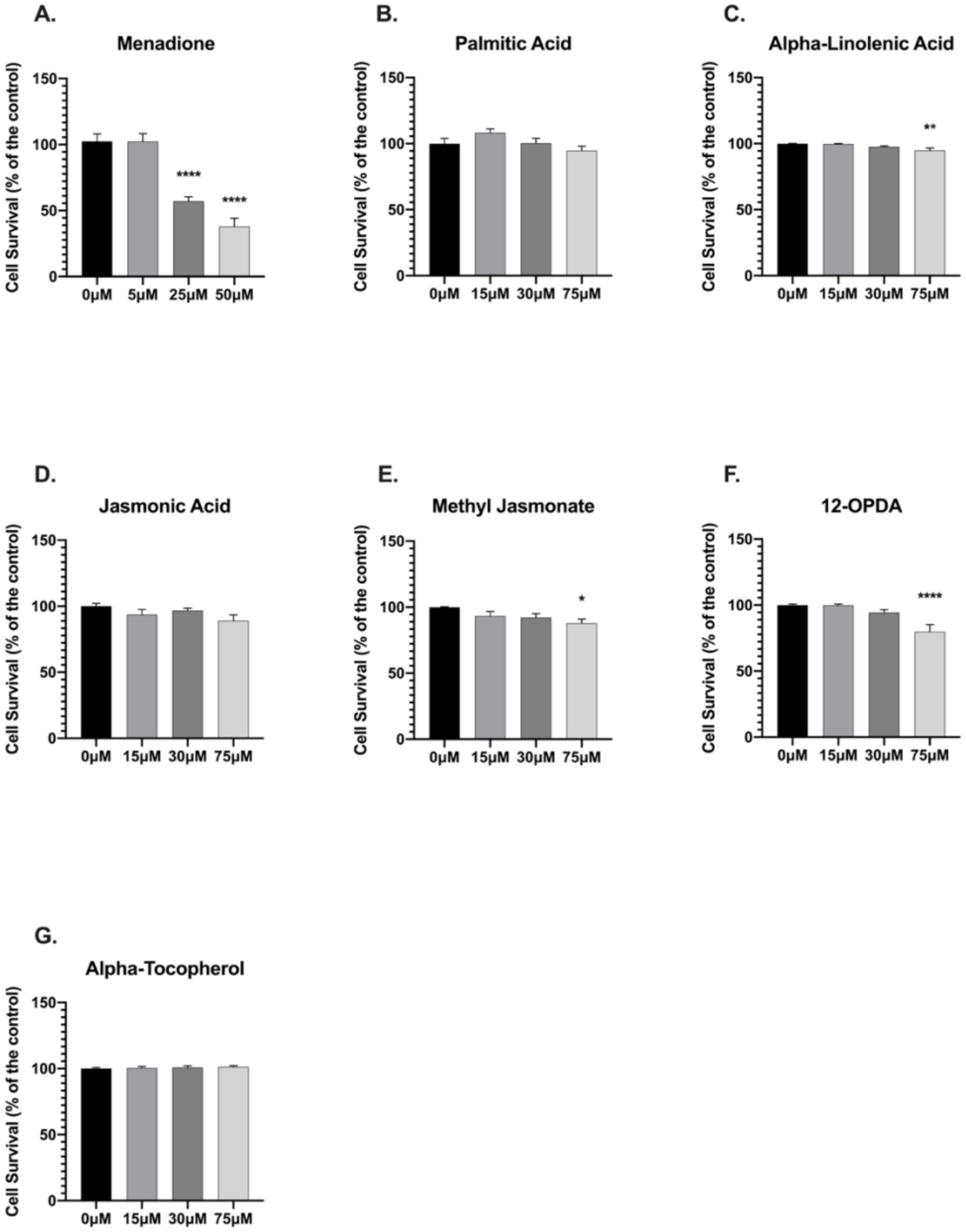

**Figure B. A superimposition of jasmonic acid and prostaglandin $E_2$** A 360° video displaying jasmonic acid superimposed over prostaglandin $E_2$ about their common cyclopentanone backbones. Prostaglandin $E_2$ is yellow and jasmonic acid is cyan. Both JA and $PGE_2$ are in the trans configuration about the five-carbon ring as was used in our study. The superimposition was performed in Jmol: an open-source Java viewer for chemical structures in 3D (http://www.jmol.org/). The close alignment can be quantified using the root mean square difference (RMSD) 1.68, or 0.02 Angstroms. The visual aid was used to support our hypothesis that the jasmonate mechanism of action may rely on structural similarities to $PGE_2$.

YouTube link: "Superimposed Prostaglandin E2 and Jamsonic Acid" https://youtu.be/AKhADwRJEAg

**Table A. Full isoform list used in PCR expression data analysis** We analyzed the PCR expression data for sixteen isoforms covering four subtypes of the E prostanoid receptor following retinoic acid treatment of SH-SY5Y cells. Included were three isoforms of *ptger1* transcripts, one of *ptger2*, nine of *ptger3*, and three of *ptger4*. The transcript isoforms of the receptors EP 1-4 were identified using their Affymetrix probe ID, or NCBI Protein accession number (*Protein, NCBI, 2004*). The full list can be found below.

| | **gene name** | **Affymetrix probe ID** (NCBI Protein accession number) |
|---|---|---|
| 1 | ptger1 | 207650_x_at |
| 2 | ptger3 | 208169_s_at |
| 3 | ptger3 | 210375_at |
| 4 | ptger3 | 210831_s_at |
| 5 | ptger3 | 210832_x_at |
| 6 | ptger3 | 210833_at |
| 7 | ptger3 | 210834_s_at |
| 8 | ptger3 | 211265_at |
| 9 | ptger3 | 213933_at |
| 10 | ptger1 | 214391_x_at |
| 11 | ptger4 | 217158_at |
| 12 | ptger3 | 231030_at |

| | | |
|---|---|---|
| 13 | ptger1 | 231201_at |
| 14 | ptger4 | 204896_s_at |
| 15 | ptger4 | 204897_at |
| 16 | ptger2 | 206631_at |

**Figure C. Normalized immunoassay results used to declare significance** The results of our enzyme immunoassays for intracellular cAMP were normalized against the relevant $PGE_2$-only treated control condition– either the 30µM $PGE_2$ or 75µM $PGE_2$-treated wells. Changes in cAMP levels were assessed for significance using a post-hoc comparison of the means of N=4 experiments for each set of conditions. Significance was established using normalized values to account for inter-test variability. The established significance levels associated with each experimental condition were transferred over to graphs representing discrete (not normalized) concentrations of cAMP (see: **Figure 6**).

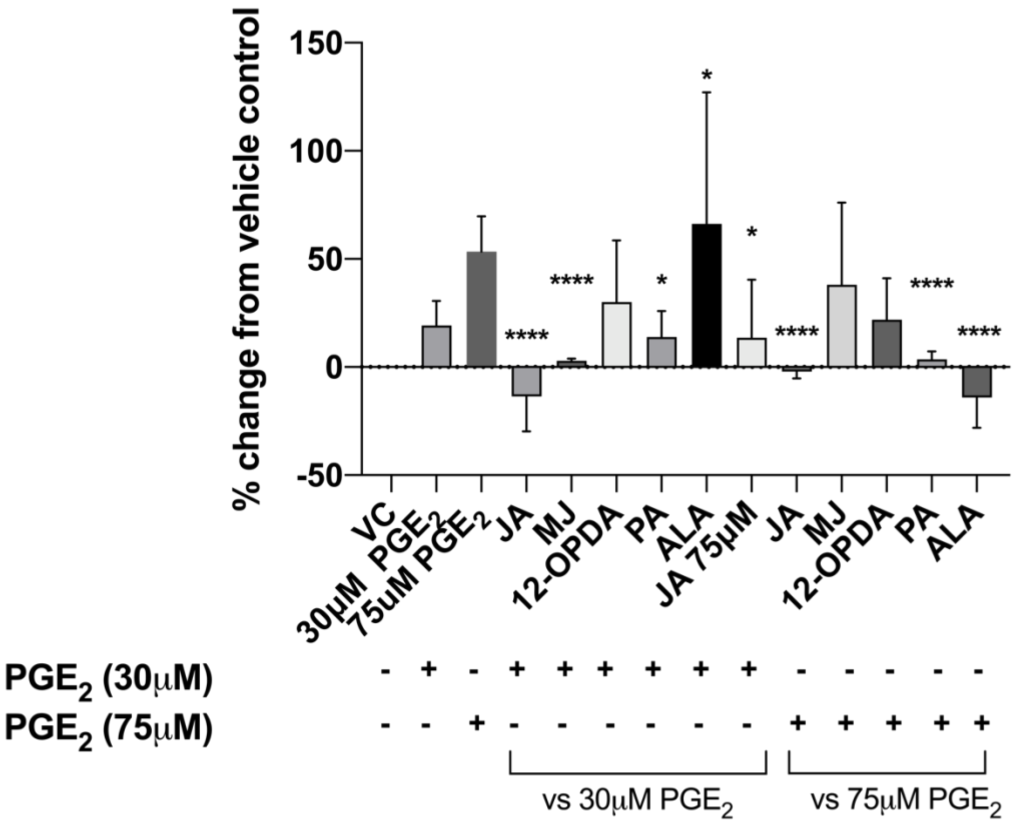
